# Adaptive Wheel Exercise for Mouse Models of Parkinson’s Disease

**DOI:** 10.1101/2024.06.19.598025

**Authors:** Henry Skelton, Dayton Grogan, Amrutha Kotlure, Ken Berglund, Claire-Anne Gutekunst, Robert Gross

## Abstract

Exercise is widely studied as a therapy in mouse models of neurological disease. However, the established techniques for exercise in mice are not ideally suited toward motor deficient disease models, nor do they facilitate active measurement of neurophysiology with tethered assays. To address this, we developed an apparatus and technique for inducing exercise in mice without aversive stimuli, using a motorized wheel with closed-loop acceleration that tracks subject performance. We demonstrated the efficacy of this approach in the 6-hydroxydopamine mouse model of PD, including with single-unit electrophysiology. This approach should allow for exercise to be better studied as a dynamic, physiological intervention in neurological disease models.

## Introduction

Physical exercise is thought to be therapeutic in a broad range of neurological diseases. Even where this is well-evidenced in patients, preclinical studies in rodents are essential to dissect the underlying mechanisms. This work is limited by the challenges of modeling physical exercise in rodents, particularly mice.

Healthy mice with access to a freely moving wheel will run spontaneously and vigorously^1^, so much so that it strains translational relevance to humans^2^. However, the tendency to exercise varies between animals^3^ and can be impaired in neurological disease models^4^. It is also challenging to perform acute assays (such as electrophysiology) on a spontaneous home-cage activity that occurs primarily in the dark cycle. Even measuring exercise performance itself is non-trivial in group-housed animals^5^.

Forced exercise offers a more controllable alternative, most commonly in the form of treadmill running motivated by an aversive stimulus. Typically a painful electrical shock, that stimulus introduces an additional variable that can differ markedly between subjects^6^, confound therapeutic effects^7–9^, and necessitate selective exclusion of poor exercisers^10^.

Motorized running wheels have been proposed as a superior, non-aversive means of inducing exercise in mice^2^ that take advantage of their natural tendency to run on wheels. This approach has been used in studies of learning and memory^11^, vascular dynamics^12^, circadian rhythm^13^ and breast cancer^14^. However, there are no reports of a motorized exercise wheel being used in a model of neurological disease. Additionally, there are some indications that the existing wheel apparatuses do not reliably induce running at the speeds typical of other modalities^13,14^.

Here, we tested a commercially available forced exercise wheel with the goal of using it in mouse models of neurological disease. Finding it unsuitable, even in healthy mice, we developed an improved platform. This comprised a physically optimized wheel with a closed-loop control system and high-quality videography for precise measurement of exercise performance. We validated this platform in the 6-hydroxydopamine (6-OHDA) model of Parkinson’s Disease (PD), including with synchronized, tethered, single-unit electrophysiology.

## Results

### Testing a Commercial Exercise Wheel

Our initial attempt at exercise used the only commercially available motorized running wheel designed for mice. Intending to apply it in the 6-OHDA mouse model of Parkinson’s Disease, we developed a longitudinal exercise regimen (figure 1A) based on a study using the same wheel^14^. Intensity was slowly increased across our 9-week intervention, for a total of 7163 meters of wheel motion, with the 6-OHDA injection surgery planned for week 5.

**Figure 1-.**
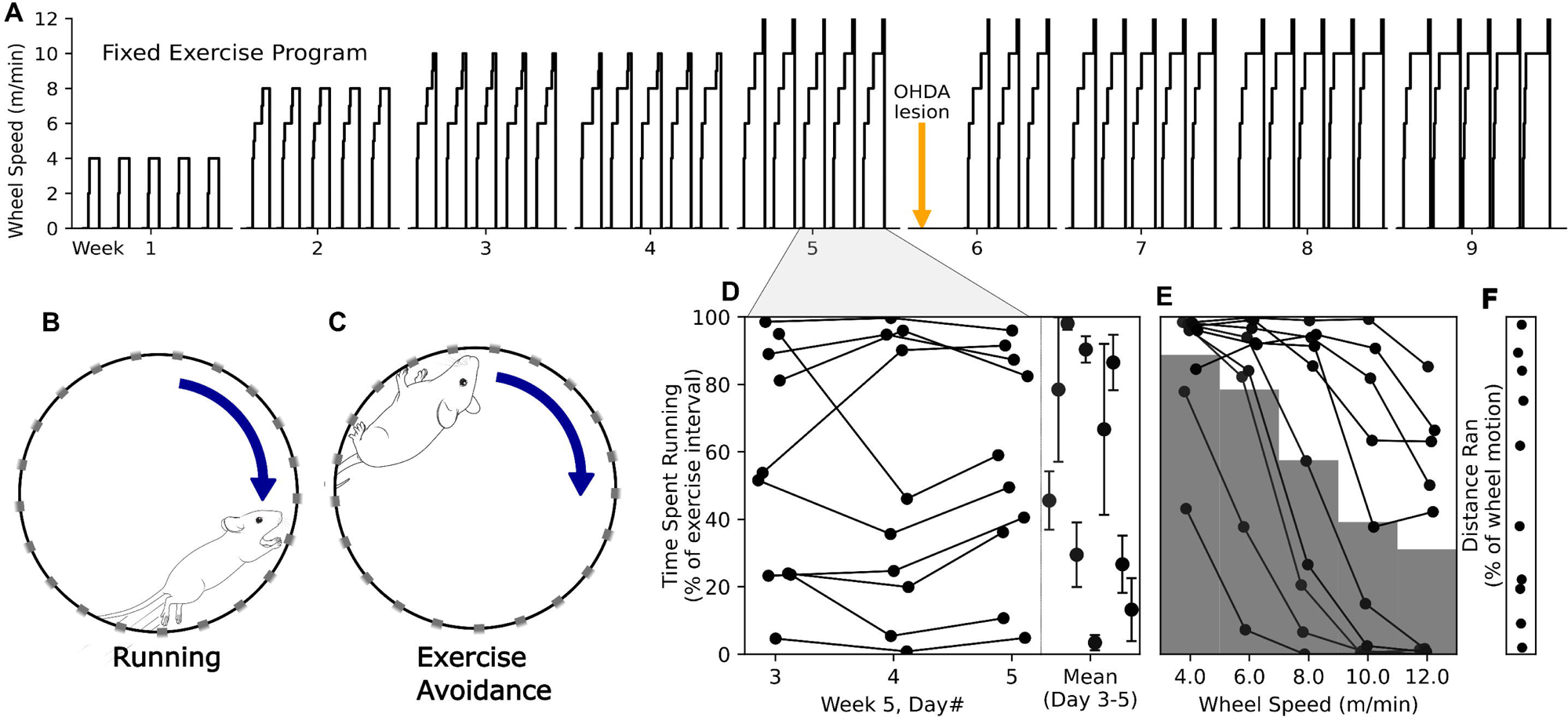
Exercise Intervention with Commercial Wheel. **(A)** The exercise program, with gradually increasing wheel speed within and across daily (5x per week) sessions. **(B)** Sketch of a mouse running versus multiple forms of exercise avoidance in **(C). (D-F)** Exercise performance scored according to the percentage of the exercise interval spent running. **(D)** Each subject’s daily exercise performance (lines) on the left, with the mean (dots) and standard deviation (bars) across days on the right. **(E)** Each subject’s performance (lines) divided according to wheel speed. **(F)** Percentage of wheel motion spent running, for each animal (dots).

Despite these efforts and even prior to the 6-OHDA lesion, we observed that rather than running (figure 1B), mice (n=10, 2 cages of age-matched C57Bl/6J males) tended to avoid exercise (figure 1C), primarily by grabbing onto the track, jumping, or sliding. Video recordings were of sufficient quality to identify these behaviors. Scoring of the video according to the percentage of exercise intervals spent running reflected a prevalent failure to exercise (figure 1D). In the final three days prior to lesioning, the percentage of time spent running ranged from 0.8 to 99.7% (mean 53.9, SD 34.8%), with a mean standard deviation of 34.5% between subjects and 10.0% across days. Running tended to decrease at higher speeds (figure 1E), and overall running distance ranged from 1.9 to 97.7% of the total wheel motion (figure 1F).

### Designing a New Wheel

As the commercial wheel was unsatisfactory, even in healthy mice, we developed an improved platform to address its shortcomings. The design is based on a much larger, non-motorized plastic wheel designed for large pet rodents (figure 2A), which does not have any surface that can be held on to. The wheel was modified for attachment to a geared motor platform (figure 2B) using a direct drive. The mouse is retained using a fully transparent front plate with a door that can admit tethered instruments, and a fixed camera for high quality, consistent videography. This design (figure 2C) can be constructed with readily available parts and simple machine work.

**Figure 2-.**
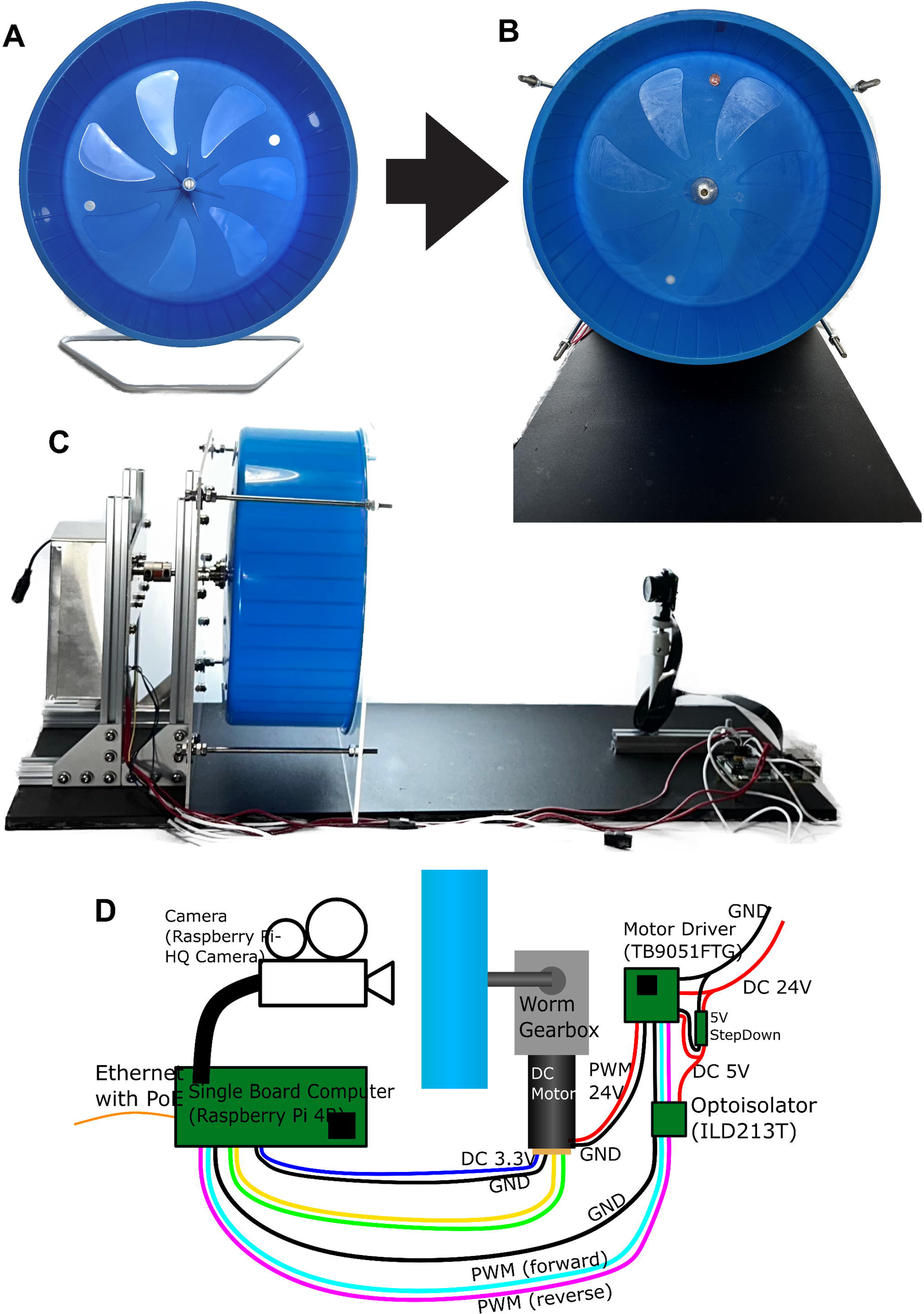
Improved Exercise Wheel Design. The free moving exercise wheel designed for large pet rodents **(A)**, shown adapted to the motorized platform in **(B**), with the modified central hub for attachment to the motor in **(C)**. **(D)** Schematic of the components and wiring for the exercise wheel.

The wheel is controlled by digital pulse-width-modulated input and outputs its position using a rotary encoder (figure 2D). This digital input, as well as the video, is controlled using customizable scripts and accessed over a local network for initiation and monitoring of experiments.

### Testing and Quantifying Exercise on the New Wheel

We tested a prototype of our new exercise wheel using the program previously used on the commercial wheel. Failure to run was much less common, and primarily consisted of sliding at the end of the wheel - mice could not hold onto the track. For comparison, we replicated the scoring of week 5. With the new wheel, running percentage ranged from 92.0 to 99.3% (mean 95.6, SD 2.4%), with a mean standard deviation of 1.3% between subjects and 2.2% across days (figure 3A). Running tended to decrease at higher speeds (figure 3B), and overall running distance ranged from 94.2 to 97.7% of the total wheel motion (figure 3C).

**Figure 3-.**
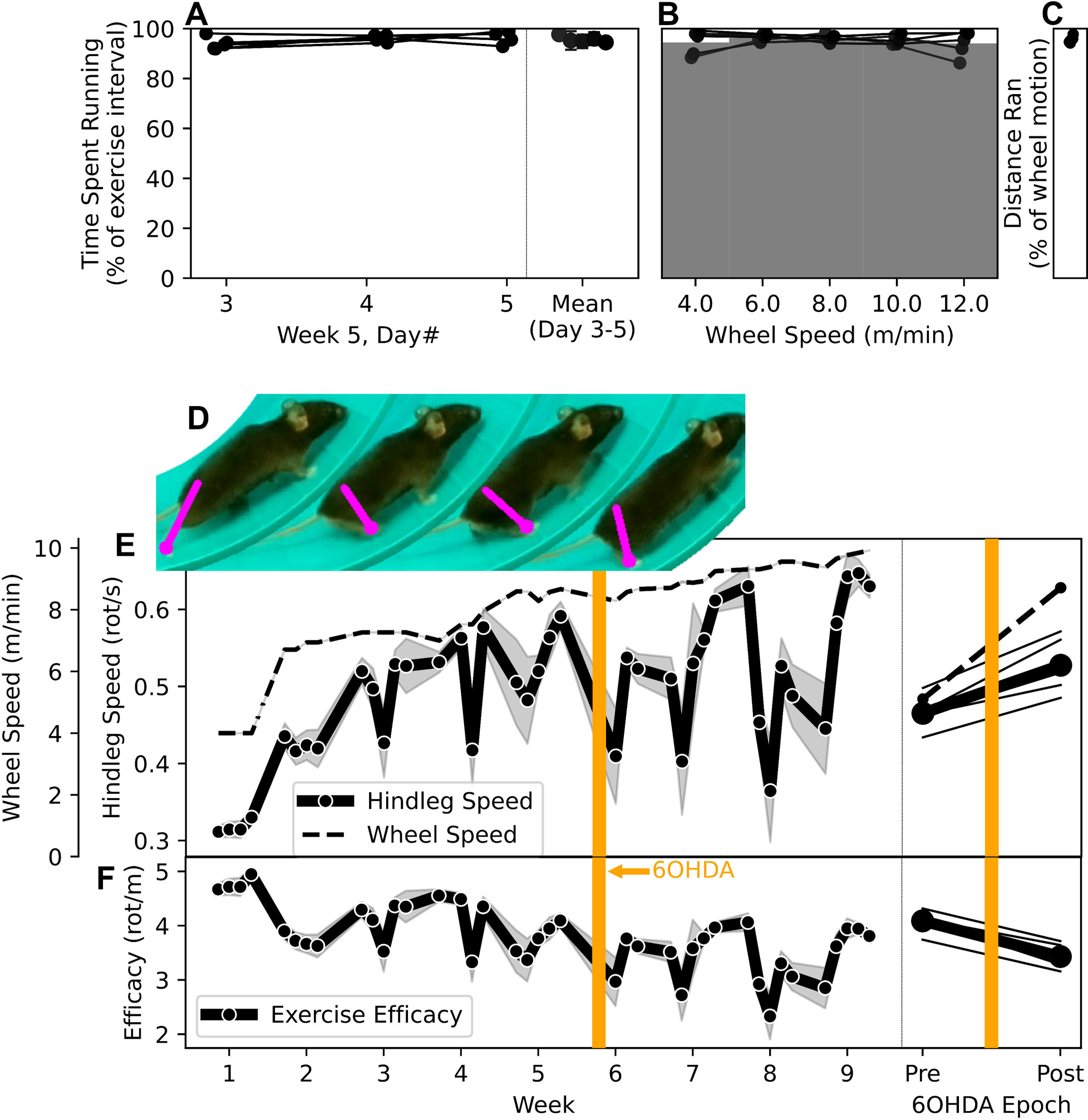
Original Exercise Regimen with New Wheel. **(A-C)** Exercise performance scored according to the percentage of the exercise interval spent running. **(A)** Each subject’s daily exercise performance (lines) on the left, with the mean (dots) and standard deviation (bars) across days on the right. **(B)** Each subject’s performance (lines) divided according to wheel speed. **(C)** Percentage of wheel motion spent running, for each animal (dots). **(D-F)** Exercise performance quantified using markerless body part tracking. **(D)** Illustration of the measurement of hindleg angle used to measure hindleg rotational speed (*w_l_*). **(E)** On the left, is the mean hindleg speed (line) and standard error (bands) for each day in magenta, alongside the wheel speed (*v_w_*) in cyan. Epoch-wise means are shown on the right. **(F)** parallels E, showing the exercise efficacy measure 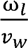.

This platform also provided an unobstructed view of the mouse, allowing for more temporally precise, objective quantification of exercise using video capture and markerless pose tracking. Taking advantage of this, we developed a measure of running based on the rotational speed of the hindleg about the hip (figure 3D). This was applied over the course of the entire intervention (figure 3E). Hindleg motion increased as the exercise program progressed in intensity, but not in proportion to wheel speed, especially after the 6-OHDA lesion. The ratio of hindleg to wheel movement, which reflects the efficacy of inducing exercise, decreased as mice became more likely to avoid exercise (figure 3F).

### Developing Adaptive Exercise

To fully capitalize on this new exercise platform, we developed an improved approach to exercise programming. Based on pilot experiments using manual control of the wheel speed with a potentiometer, we devised a strategy to maximize wheel running. Starting with the wheel at a low speed, as soon as mice started advancing along it, even newly acclimated mice could keep pace with rapid acceleration to the top speed (13.3 m/min). However, at a constant speed they would inevitably fall off-pace and start sliding, only being able to recover if the wheel was decelerated. With sufficiently rapid deceleration, mice would usually recover without failing to run entirely.

This heuristic was translated into a formal control strategy for maximizing running. The input signal was the mouse position on the wheel, which could be measured at low latency (<10ms) and with minimal computational resources using color thresholding (figure 4A). The control strategy itself was divided into three regimes, depending on the position, 0, of the mouse, with 0 = 0 at the center of the bottom of the wheel and increasing against the direction of wheel motion (figure 4B). Toward the back of the running area, where 90° < 0 < 30°, the wheel is decelerated at a rate, 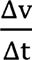, proportional to the position of the mouse, 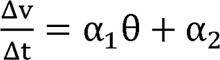. Toward the front, acceleration is proportional to the mouse position and the change in position, 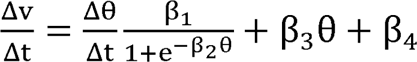. For |θ| > 90°, the program shuts off the wheel in the unlikely case that mice are able to hold onto it. Outside of this regime, the speed is limited to 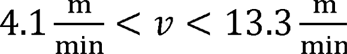, based on the limits of the gearbox and motor. The parameters, α_n_ and β_n_, were optimized empirically via pilot experiments in healthy mice.

**Figure 4-.**
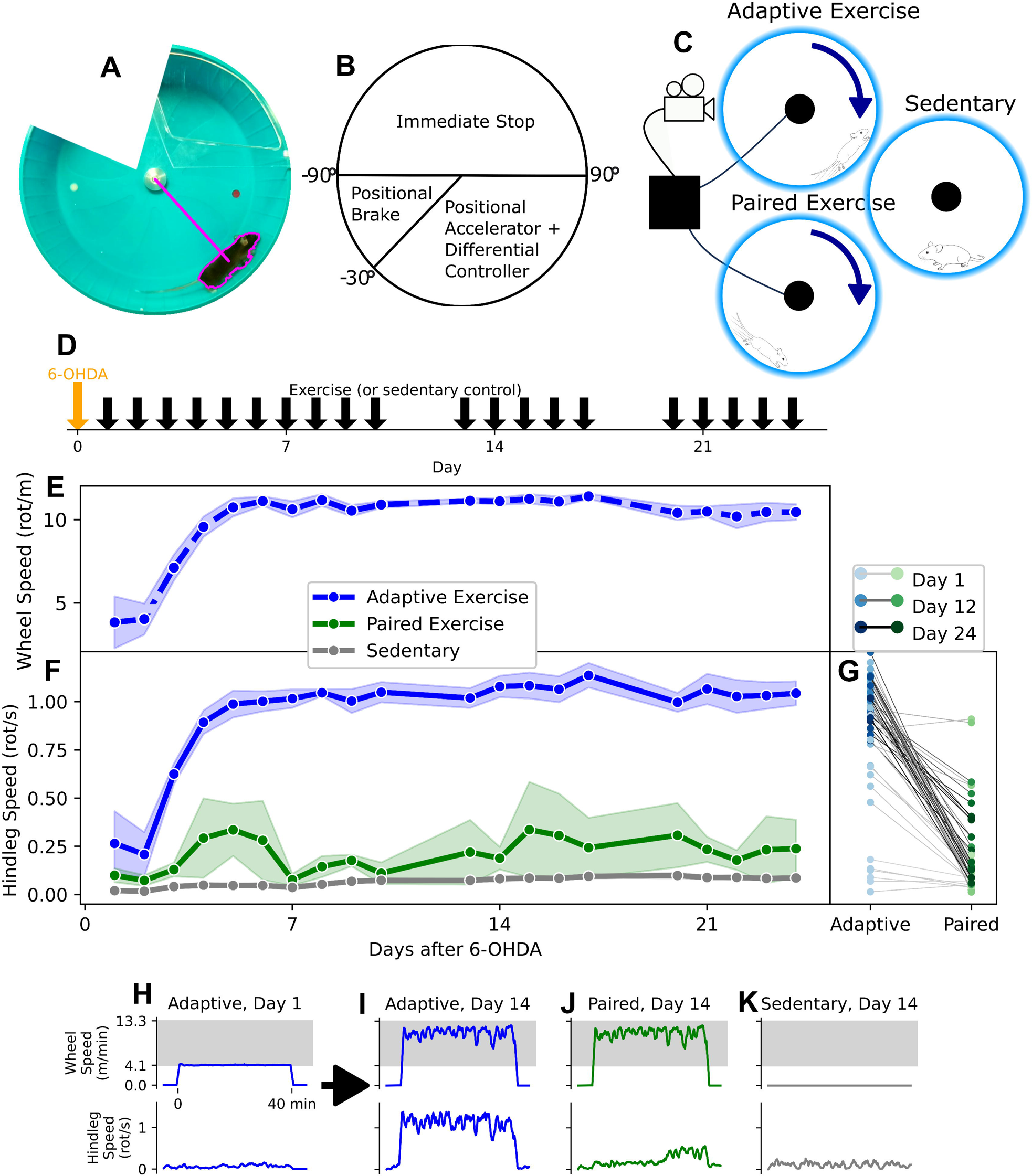
Adaptive Exercise Intervention. **(A)** An example frame of the real-time tracking, based on color thresholding, used for adaptive exercise. **(B)** Control regimes for adaptive exercise, determined according to mouse position. **(C)** Treatment groups being compared to adaptive exercise in 6-OHDA mice. **(D)** Timeline of the exercise intervention. **(E)** Wheel speed over the course of the adaptive exercise intervention, including the daily mean (dots) and standard error (bands) for each day. **(F)** Hindleg speed over the course of the intervention, including the daily mean (dots) and standard error (bands) for each day. **(G)** The daily performance for each subject (dots) in the adaptive and paired exercise pairs, with the day indicated by shading. **(H-K)** Examples of daily exercise sessions showing the wheel speed and hindleg speed.

We tested this program in disease model mice, beginning on the day after the injection procedure. We compared a group undergoing adaptive exercise (n=5) against two control groups (figure 4C). In the “Paired” group (n=4), each mouse was matched to one of the mice undergoing adaptive running and exercised using identical motor inputs. The sedentary control mice (n=5) were placed in a stationary wheel for the same length of time. This experiment was carried out over four weeks, with daily 40-minute sessions (figure 4D). There was no attrition among the adaptively exercised mice, compared to 2 (50%) of the paired mice and 2 (40%) of the sedentary mice which died due to 6-OHDA-related complications.

The wheel speed in the adaptive group was driven by animal behavior, and the mean daily speed rapidly rose and stabilized after the early effects of intrastriatal 6-OHDA resolved and animals adapted to exercise (figure 4E).

This was well-reflected in the measured hindleg motion of the adaptively trained mice, but not in the paired mice, despite identical wheel inputs (figure 4F). Among sedentary mice, whose hindleg motion reflects free movement in the stationary wheel, the mean was 0.06 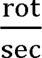 (SD 0.03). Compared to the sedentary controls, adaptive training was associated with 0.84 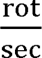 greater hindleg motion, a significant difference (p=6.1×10^−43^), while paired training was associated with a non-significant 0.12 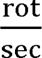 increase (p=0.061). Directly comparing the adaptively trained mice and their paired counterparts in a pairwise fashion, adaptive training was associated with 0.65 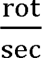 greater hindleg motion (p=2.5×10^−14^).

In the early adaptive exercise sessions, when mice failed to engage with exercise (primarily by sliding), the wheel speed remained near the minimum speed (figure 4H). As they began to run with the wheel, the control program produced a bursty, high-speed interval pattern, which was reflected in hindleg speed (figure 4I). At the same point in training, paired mice had identical wheel movement, without similar hindleg movement (figure 4J). Sedentary mice, with a stationary wheel, had minimal, steady hindleg motion (figure 4K).

To quantify the evolution of adaptive training over time, the average daily wheel speed for each mouse was fit to a logistic function, 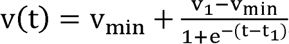, with an asymptote (*v*_1_) at the stabilized wheel speed, and an inflection point (*t*_1_) where adaptation to exercise occurred. The mean *v*_1_ speed was 10.9 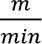 (SD 0.53), and the mean *t*_1_ was at 2.9 days (SD 1.29). Similarly, each adaptively trained mouse’s performance was measured by fitting to a function, 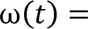 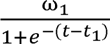. The mean stable hindleg speed (ω_1_) was 1.05 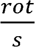 (SD 0.11), and the mean *t*_1_ was at 2.5 days (SD 1.3).

### Demonstrating Tethered Electrophysiology Adaptive Exercise

Finally, we demonstrated that adaptive wheel exercise could be used in conjunction with electrophysiology, as an example of a tethered, noise-prone assay. The wheel was modified with a Faraday cage, a higher-speed camera with shutter signal output, and a computer-based control and recording platform (figure 5A). Electrophysiology and all digital signals were recorded using the same clock for precise synchronization (figure 5B).

**Figure 5-.**
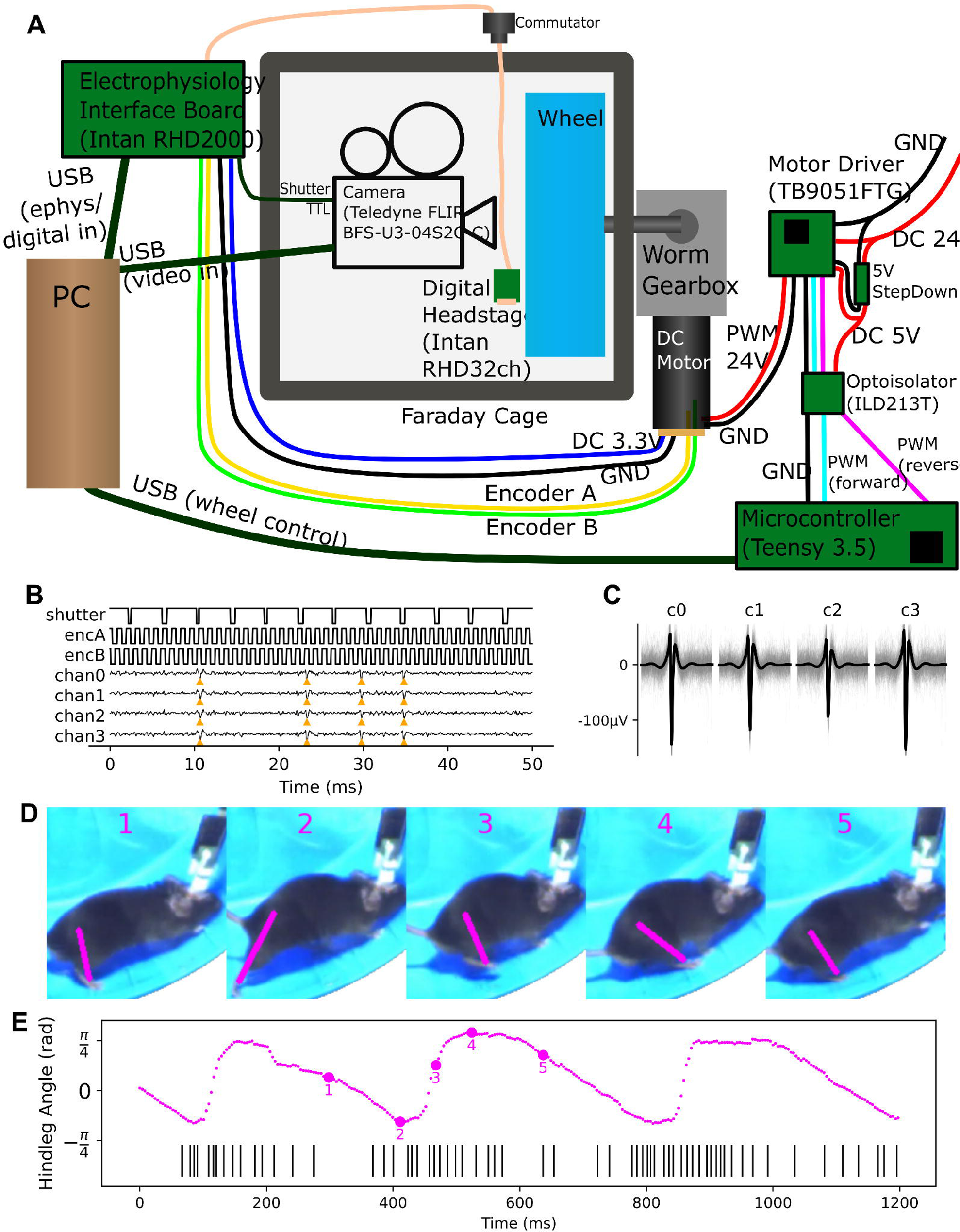
Adaptive Exercise with Electrophysiology. **(A)** Circuit diagram of the running wheel exercise apparatus, modified for synchronized electrophysiology and high-speed videography. **(B)** An example of recorded data, showing the digital signals from the camera shutter and the wheel’s rotatory encoder, captured in precise synchronization with electrophysiology. Below, the four analog traces are shown after high-pass filtering and common average referencing, with neural spikes from a sorted single unit noted (orange triangles). **(C)** A subset of waveforms from the unit noted in **B**, along with the overall mean waveform. **(D)** Frames from a short (approximately 500ms) segment of video, showing the mouse running along the wheel, with the identified hindleg angle indicated (magenta line). **(E)** A plot showing the hindleg angle over the course of three periods of the gait cycle alongside spikes from the unit in **C**, with the frames in **D** indicated (magenta numbers).

Using the adaptive exercise program, after several sessions of unencumbered training, tethered mice are consistently able to run at high speed. This has been confirmed in normal and 6-OHDA lesioned mice (n=5) implanted with multi-tetrode microdrives, and allows for the precise measurement of correlates between neural activity and behavior during exercise. As an example, the spiking of a single unit in the substantia nigra (figure 5C) was tracked alongside the gait cycle (figure 5D) as a 6-OHDA-lesioned mouse ran along the wheel for 1.2 seconds (figure 5E).

## Discussion

Despite theoretical advantages to motorized wheel exercise, existing approaches fall short in being able to consistently induce running in mice, even absent a motor deficit. Here, we developed an improved wheel and adaptive exercise paradigm and demonstrated its efficacy in the 6-OHDA model of Parkinson’s Disease. This should provide a non-aversive means of inducing running in mouse models of neurological disease that is amenable to precise quantification and tethered assays.

Even using a low-intensity training program with gradual speed increases, the commercially available exercise wheel did not consistently induce running. The actual behavior of treated mice varied greatly, with half rarely or never running at the highest speeds. Despite identical treatment, the subjects did not undergo comparable interventions, and for many it was not truly a running exercise intervention at all. Moreover, due to the limited wheel space and the manner of failure to exercise, it was unfeasible to carry out tethered experiments.

This is not necessarily unexpected. Of the four reports that describe motorized wheel exercise in mice, only two indicated that they verified running performance. One of those specifically describes the motorized wheel as unsuitable at higher speeds^13^. The other^14^ reports it as being poorly suited to pre-defined exercise programs. Instead, they used a custom motor system that allowed the experimenter to actively control speed in response to mouse behavior, increasing it very carefully within the limits of the poorest performing mouse in a cohort. That approach was not used here, due to the difficulty of replicating manual coaching.

Our improved exercise wheel provided for much more consistent running. Using the original exercise regimen, and scored at the same pre-6-OHDA time-points, all mice consistently ran for almost all of the exercise interval at all speeds. This was likely accomplished by a combination of the track being much larger, which facilitates running, and having fewer graspable surfaces, which averts freezing. Mice are known to voluntarily run more on larger, non-motorized wheels^15^, and freezing is a well-described pattern of exercise avoidance in motorized wheel exercise^13,14^.

In addition to an improved running track, our design facilitates high quality videography, allowing us to objectively and precisely measure running according to leg movement. On the fixed exercise program, running tended to increase with wheel speed, but they were not perfectly coupled, especially in the post-6-OHDA period. This was true in spite of a low intensity regimen with a prolonged period of acclimation, which were necessary compromises to accommodate the original wheel. It was clearly not possible to truly *force* exercise with a motorized wheel.

With that in mind, our adaptive exercise program replaces the fixed exercise program with a control strategy that responds to behavioral cues. This allowed us to effectively exercise mice in the acute phase of a PD-model lesion, without any previous training. In practice, the adaptive exercise program provided for an initial slow period of exercise acclimation and accommodated the natural tendency of mice to run in brief, high intensity bursts. However, the adaptive aspect itself was critical, as the paired mice on the same program as adaptively trained counterparts greatly under-performed them. Whilm e the adaptive program does create intersubject differences in exercise programming, this may not be a true disadvantage, as identical programming does not provide for consistent behavioral engagement.

The simplicity and efficacy of the 6-OHDA lesion, in combination with the plethora of transgenic lines available in mice, have proven particularly useful in dissecting the neurophysiological underpinnings of therapies^16–18^. Unilateral lesioning affords the additional benefit of lateralized behavioral and histologic measures. With that in mind, our chosen model may be ideally suited to probe the dynamic neural effects of exercise on the dopamine depleted basal ganglia. However, this work is among the first reports of exercise in 6-OHDA-lesioned mice, and the only one with intervention in the therapeutically critical^19^, earliest period of the disease process. That may be because this model, particularly in that time period, imposes severe motor and non-motor impairment. Mice debilitated by the acute effects of 6-OHDA may be unable or disinclined to run. There are only three other reports of exercise after a 6-OHDA lesion in mice. One used treadmill exercise starting seven days after the lesion, without mention of running performance, attrition, or the use of aversive stimuli^20^. Another used forced swimming exercise starting at the same, delayed timepoint^21^. A final study used voluntary exercise after four weeks, and only alongside treatment with levodopa^22^.

Motorized wheel running is more established in rats^23^, and there is a comparable line of work improving wheel design and programming for models of neurological disease^24,25^. That design does not allow for a tether or unobstructed video recording, but does incorporate positional tracking using infrared sensors. There, position alone is used to grade exercise and to enable a more limited responsive exercise approach^26^. In general, the construction and hardware configuration are much more complex than our design, and do not allow for measurements other than position.

Finally, beyond just invoking running, adaptive wheel exercise also allowed for the measurement of neurophysiology during exercise. This has not previously been described during rodent exercise interventions, despite the essential role of dynamic physiologic measurements in characterizing neurological disease and therapies. This is particularly relevant in a Parkinson’s Disease model, given the well described pathophysiology of that disease, and the link between investigations of pathophysiology and the development of circuit therapies. Additionally, the ability to “break” exercise, as in the ineffective “Paired” program, is important in being able to test manipulations that preclude exercise (and the associated physiology). This would not be feasible on a platform that uses aversive therapy to force exercise.

This work is limited by testing our exercise intervention in a single disease model and assessing it solely according to animal movement. Future work should expand this work to other models of Parkinson’s Disease as well as models of other neurological diseases. Additionally, this intervention should be further validated according to additional measures and potential confounders of exercise performance, including cardiovascular and metabolic indicators as well as stress markers.

## Conclusions

Adaptive wheel exercise, including both the improved physical apparatus and closed-loop controller described here, provides for effective, non-aversive induction of running in a mouse model of late-stage Parkinson’s Disease. This approach should enable a better understanding of the effects of exercise in disease models with motor deficits, particularly the acute neural effects, without the confounds imposed by aversive stimuli.

## Methods

### Animals

All animal studies were carried out in accordance with animal-use protocols approved by the Emory University Institutional Emory University Institutional Animal Care and Use Committee. Male C57Bl/6J strain mice (The Jackson Laboratory, strain# 000664) between 8 and 12 weeks of age were used for all experiments.

### Parkinson’s Disease Model

Hemiparkinsonism was induced by unilateral intrastriatal injection of 6-OHDA. Placed on a stereotactic frame (Kopf Instruments) under isoflurane anesthesia, a craniostomy was created at 1.8mm lateral and 0.4mm anterior to bregma, and a glass needle lowered 3.25mm ventral from the dural surface. At this site, a microinjector (Nanoject 3, Drummond Scientific) was used to inject 1µL of 6-OHDA hydrobromide (Sigma Aldrich #162957) solution (2.67 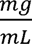, in 0.9% saline + 0.2% ascorbic acid). For two weeks following surgery, animals were weighed daily and provided supplementary high-calorie food (ClearH_2_O #72-04-5022) and intraperitoneal fluids as needed.

### Commercial Motorized Exercise Wheel

Motorized exercise wheels were sourced from Lafayette Instruments (Model 80840B), the only vendor with a model designed for mice. These were controlled using the included Scurry software. The wheels were modified with the addition of rubber tracks and with holes on the sidewall blocked to limit grabbing onto the wheel. Exercise was recorded using an external video camera.

### Fixed Exercise Programming

The exercise regimen for the commercial wheel and initial testing of the new design consisted of daily (5x/week), 20-minute exercise sessions on the moving wheel. Each session included fixed periods of time at speeds that gradually increased within each session and across sessions. These speed settings were based on a study where the same wheel (using a different, customized motor platform) was controlled by an experimenter to not exceed the perceived running capacity of the poorest runner among a large cohort^14^. The 9-week exercise treatment was interrupted at week 5 with the surgical 6-OHDA lesion, after which point the mouse was allowed two days without exercise to recover.

### Manual Exercise Scoring

Exercise videos were manually scored using BORIS^27^ to mark the beginning and end of running intervals, which were defined as periods where the mouse was running along the wheel, rather than holding onto it, sliding, or jumping. Each exercise interval was assessed according to the percentage of total time spent running.

### Custom Wheel Construction

The driveshaft, backplate, frontplate, motor plate, base, and support rails were cut and drilled to shape. The original wheel (Kaytee #045125613827) was modified by machined away the original hub and replacing it with a custom aluminum hub to attach to the drive shaft flange, with the original wheel axle and stand discarded. From there, all parts were assembled without further modification.

### Wheel Programming

The motor is controlled via pulse width modulated input to the motor controller, and wheel position is read out with a quadrature rotary encoder. Both are connected to general purpose digital input/output pins on a single-board computer (Raspberry Pi 4B, Raspberry Pi Foundation). The camera (High Quality Camera, Raspberry Pi Foundation) is connected to that device for integrated camera input. All systems are controlled using custom Python scripts with Jupyter notebooks (accessed over a local network) as the interface to run experiments. Exercise programs (including the adaptive controller, described below) automatically record video (90fps) and for each frame save a count of edge detections from the rotary encoder.

### Automated Exercise Quantification

Pose detection with DeepLabCut^28^ was used to identify key points along the mouse body for each frame of video. The coordinates of the hip joint and the hind foot on the visible side when the mouse faced forward on the wheel were used to calculate a hindleg angle. The absolute difference in that angle between subsequent frames was calculated as the leg movement, which was divided by the assessed time periods to give the leg speed. This was reported as full rotations (2π radian) per second.

Wheel motion was determined using the count of rotary encoder edge detections saved for each frame of video. The track motion was calculated based on the resolution of the encoder, gearing, and circumference of the track. Exercise efficacy was calculated as the ratio of leg movement to wheel movement.

### Adaptive Exercise Programming

The adaptive exercise controller uses a live sampling of the recorded video to determine mouse position. Prior to each experiment, the wheel center coordinate and color settings for segmentation of the mouse against the blue wheel background can be adjusted. For each frame, the largest color-thresholded region is identified as the mouse, with the angle between the wheel center and the mouse centroid calculated as the mouse location.

### Adaptive Exercise Comparison

To validate the adaptive controller in 6-OHDA mice, exercise was applied in daily, 40-minute sessions for 4 weeks starting one day after 6-OHDA injection. The motor commands used for the adaptively animals were saved and used to run the same program as a fixed script for their paired counterparts. Sedentary animals were placed in a locked wheel for the same period of time.

### Electrophysiology Configuration

In order to perform electrophysiologic recording during adaptive exercise, the wheel system was reconfigured to be run from a computer, with all digital signals routed through the electrophysiology interface board (RHD USB Interface Board, Intan). The computer controlled the wheel via serial input from a microcontroller (PJRC Teensy 3.5), recording video using a camera connected with USB (Teledyne FLIR). Recordings and wheel control were coordinated by a custom workflow in the Bonsai^29^ software package.

### Electrophysiology Experiments

To prepare mice for electrophysiologic recording, two weeks after 6-OHDA surgery, they were implanted with custom-fabricated micro-drive tetrode arrays at 1.4mm lateral and 3.2mm posterior to bregma. One week was allowed for recovery, and animals were acclimated to adaptive exercise over 3-5 sessions before beginning tethered experiments. These experiments used an identical adaptive control strategy, with exercise similarly applied over 40 minutes sessions. In brief, electrophysiologic recordings were preprocessed using a bandpass filter (600 to 6000 Hz), followed by common median referencing, and spike sorting with Kilosort 2^30^. Single units were identified according to the designation provided by Kilosort in addition to an interspike interval violation ratio less than 0.1 and a signal-to-noise ratio greater than 4.

### Statistics

Hindleg movement was compared between groups via linear mixed-effect models, with fixed effects associated with treatment group and interventional day and subject level random effects. Pairwise comparison between the adaptively trained mice and their matched counterparts included the shared exercise program as an additional random factor.

## Acknowledgements

This work was supported by funding from the Department of Defense Congressionally Directed Medical Research Programs (Award no. W81XWH-19-1-0776).

